# CD42b immunostaining as a marker for placental fibrinoid in normal pregnancy and complications

**DOI:** 10.1101/860973

**Authors:** Peilin Zhang

## Abstract

**Background:** There are two types of the fibrinoid deposits within the placenta, and the nature of these fibrinoid are poorly understood in clinical setting.

**Design:** Fibrinoid deposits within normal pregnancy and the pregnancy related complications are studied using the routine hematoxylin and eosin stain and immunostaining for CD42b as a marker for platelet aggregates and fibrin.

**Results:** Fibrin-like fibrinoid is associated with platelet aggregates characterized by the positive immunostaining for CD42b and coagulatory cascade activation with blood flow changes in the circulation and intervillous spaces. Matrix-type fibrinoid is not associated with platelet aggregates and coagulation and its pathogenesis is unknown.

**Conclusion:** Fibrinoid deposits within the intervillous spaces are mostly from maternal circulation and these fibrinoids are likely the result of the laminar blood flow change at specific anatomic locations, leading to activation of coagulatory cascade. The pathogenesis of matrix-like fibrinoid is unclear. CD42b immunostaining is helpful in difficult cases.

## Introduction

Fibrinoid deposit is commonly present on the placental examination in normal and abnormal pregnancies, and it is associated with maternal and fetal complications [1]. Fibrinoid deposits are generally classified as two categories by their biochemical characteristics, fibrin-type fibrinoid, and matrix-type fibrinoid [2, 3]. Fibrin-like fibrinoid is considered to be a component of maternal coagulatory cascade leading to the traditional thrombosis, whereas matrix-type fibrinoid is likely the product of trophoblasts secretion and cellular interaction in response to a variety of molecular signals [2]. In the fetal and maternal vessels, the term “intramural fibrin deposition” was recommended by the recent guideline for maternal or fetal vascular thrombosis, and no specific designations of other types of fibrinoids were recommended in the guideline [4]. The fibrinoid materials within the intervillous spaces including classically Langhan’s fibrinoid (Langhan’s stria, subchorionic fibrinoid), Rohr’s fibrinoid (Rohr’s stria, basal plate fibrinoid), and intervillous thrombi are considered to be fibrin-like fibrinoid; whereas in maternal floor infarction (massive peri-villous fibrinoid deposits, MFI/MPFD), a critical placental abnormality with significant fetal consequences, is considered to be the matrix-type fibrinoid [1, 2, 5]. All these fibrinoid deposits are traditionally referred to as “fibrinoid” instead of “fibrin” due to the fact that the nature of these homogenous eosinophilic materials under light microscope using the routine hematoxylin and eosin stain is uncertain. Fibrinoid material in general is believed to play physiological roles in placental and fetal development, but different fibrinoid deposits in various anatomic sites are unlikely to be identical due to the geographical locations alone [2, 5]. To distinguish and specify these fibrinoids according to their anatomic locations appears to be helpful in understanding its pathophysiology and its related maternal and fetal complications. During placental examination it is generally not an issue for a pathologist to distinguish the intervillous thrombosis at the various locations and placental infarcts from maternal floor infarction (massive peri-villous fibrinoid deposit, MFI/MPFD) except for rare difficult cases [6, 7]. Some degrees of various combinations of the two types of fibrinoids can also be seen. In current study, I set to use immunostaining for CD42b to identify the platelet aggregates and its associated fibrin deposit within the placental tissues in difficult cases to distinguish the fibrin-like fibrinoid from the matrix-type fibrinoid. CD42b is also known as platelet glycoprotein 1b alpha chain (GP1BA), which is a component of the heterodimeric structure that functions as a receptor for von Willibrand factor (VWF) [8]. CD42b is expressed on the platelets and the megakaryocytes, and mutations of GP1BA gene are clinically associated with Bernard-Soulier syndromes and platelet-type von Willibrand disease [9-11]. CD42b complex is also a target for a variety of anti-thrombotic drugs in clinical use or in development [12]. The practical utility of using the CD42b immunostaining in the current study is to help in diagnosis of MFI/MPFD from its mimics with implication of its pathogenesis, since the diagnosis of MFI/MPFD is significant with implications of recurrence and fetal demise [13].

## Materials and methods

The study is exempt from Institutional Review Board (IRB) approval according to section 46.101(b) of 45CFR 46 which states that research involving the study of existing pathological and diagnostic specimens in such a manner that subjects cannot be identified is exempt from the Department of Health and Human Services Protection of Human Research Subjects. Placentas submitted for pathology examination for a variety of clinical indications are included in the study, and when the fibrinoid material was found on the routine hematoxylin and eosin stain slides, immunohistochemical staining for CD42b was performed. Placentas from normal pregnancies without clinical or pathological indications are not submitted for pathology examination in our institution.

Paraffin embedded tissues from routine surgical pathology specimens and routine hematoxylin and eosin (H & E) stained pathology slides were examined using light microscopy using the Amsterdam criteria [4]. No special procedures or stains were employed. Immunohistochemical staining procedures were performed on paraffin embedded tissues using Leica Biosystems Bond III automated immunostaining system following the manufacturing instruction. CD42b monoclonal antibody was purchased for clinical in vitro diagnostics from ABCAM (mouse monoclonal antibody against human, Catalog#: ab134087) with appropriate dilutions and controls.

## Results

The subchorionic fibrinoid (Langhans ‘stria, Langhan’s fibrinoid layer), and the basal fibrinoid (Rohr’s stria) are consistently associated with platelet aggregates that are positive for CD42b signals by immunohistochemical staining (Figure 1 and 2). The platelet aggregates and CD42b signals appears to be on the surface of fibrin material facing the intervillous spaces, not in the deep fibrin close to the chorionic tissue or basal plate tissue. Peri-villous fibrinoid within the intervillous thrombi is also found to be associated with platelet aggregates, and these can be identified in placental lesions such as accelerated villous maturation in preeclampsia, placental infarcts and infectious villitis (Figure 3 and 4). Maternal floor infarction (massive peri-villous fibrinoid deposition, MFI/MPFD) with intravillous/perivillous fibrinoid deposits is not associated with platelet aggregation and/or CD42b signals, and the etiology of MFI/MPFD was not related to maternal coagulopathy or coagulatory cascade activation (Figure 5). The trophoblastic cyst content (not shown, X-cell cyst) and the fibrinoid medial necrosis of the decidual vasculopathy are not associated with platelet aggregates or increased CD42b signals (Figure 6). There is no practical significance to identify the Nitabuch fibrinoid on the decidual surface during placental examination, as Nitabush fibrinoid connects the decidua to the uterine wall (myometrium).

**Figure 1:**
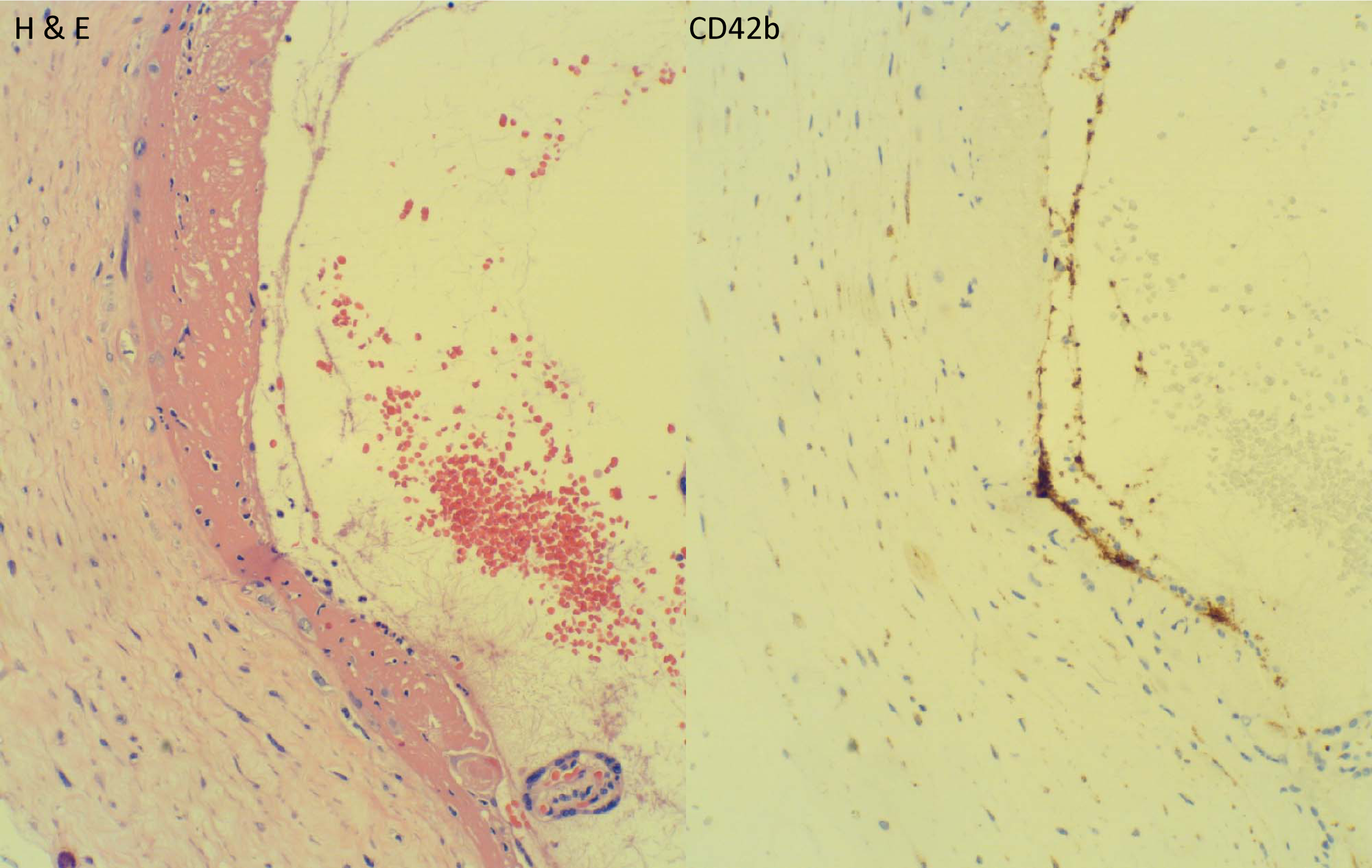
Langhan fibrinoid. Subchorionic fibrinoid (Langhan’s fibrinoid, Langhhan’s stria) with H & E stain and the corresponding immunohistochemical staining for CD42b. The platelet aggregates are highlighted by the brown immunoperoxidase signals. Unless otherwise stated, all photographs were taken at 200X magnification.

**Figure 2:**
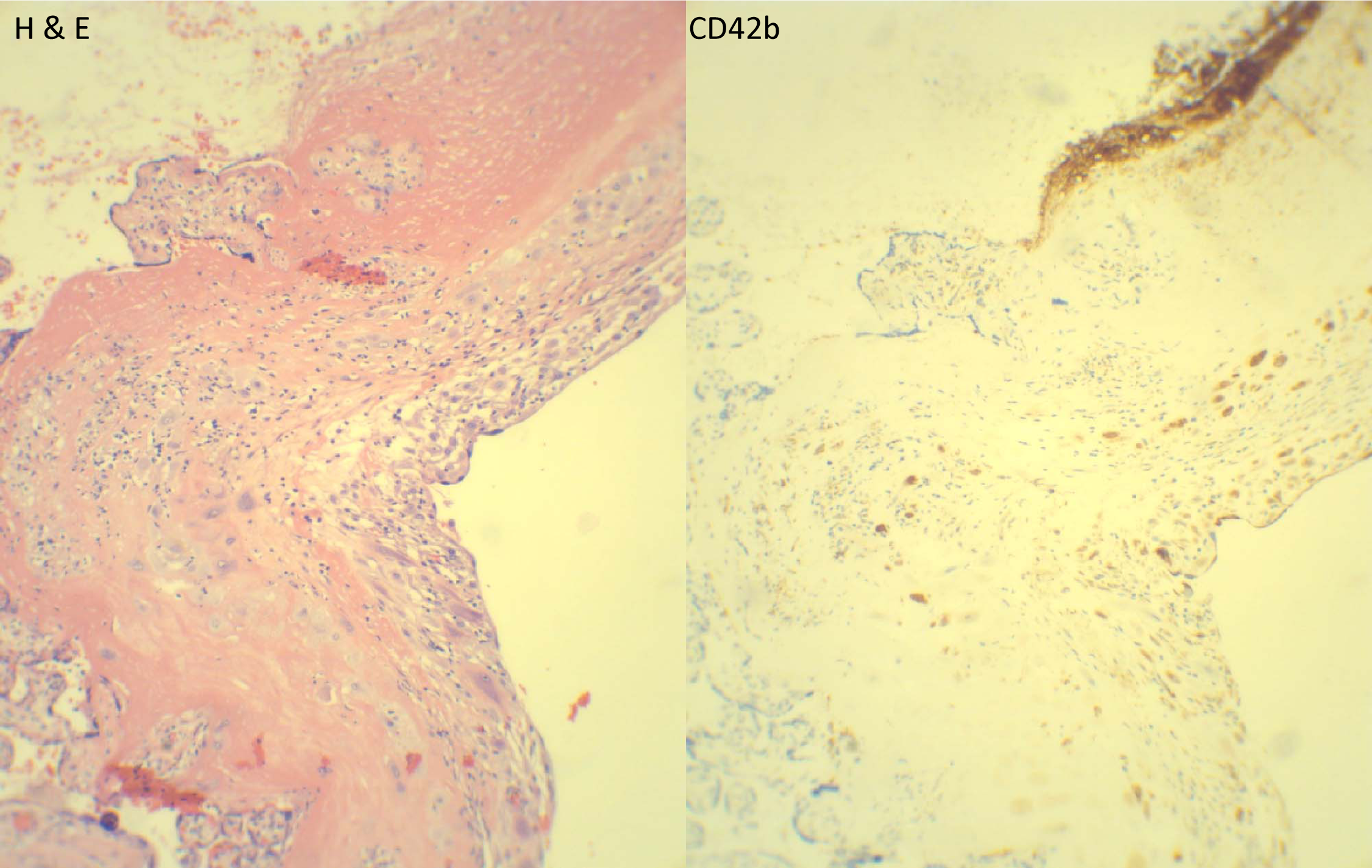
Rohr’s Fibrinoid. Basal fibrinoid (Rohr’s fibrinoid, Rohr’s stria) with H & E stain and the corresponding immunohistochemical staining for CD42b.

**Figure 3:**
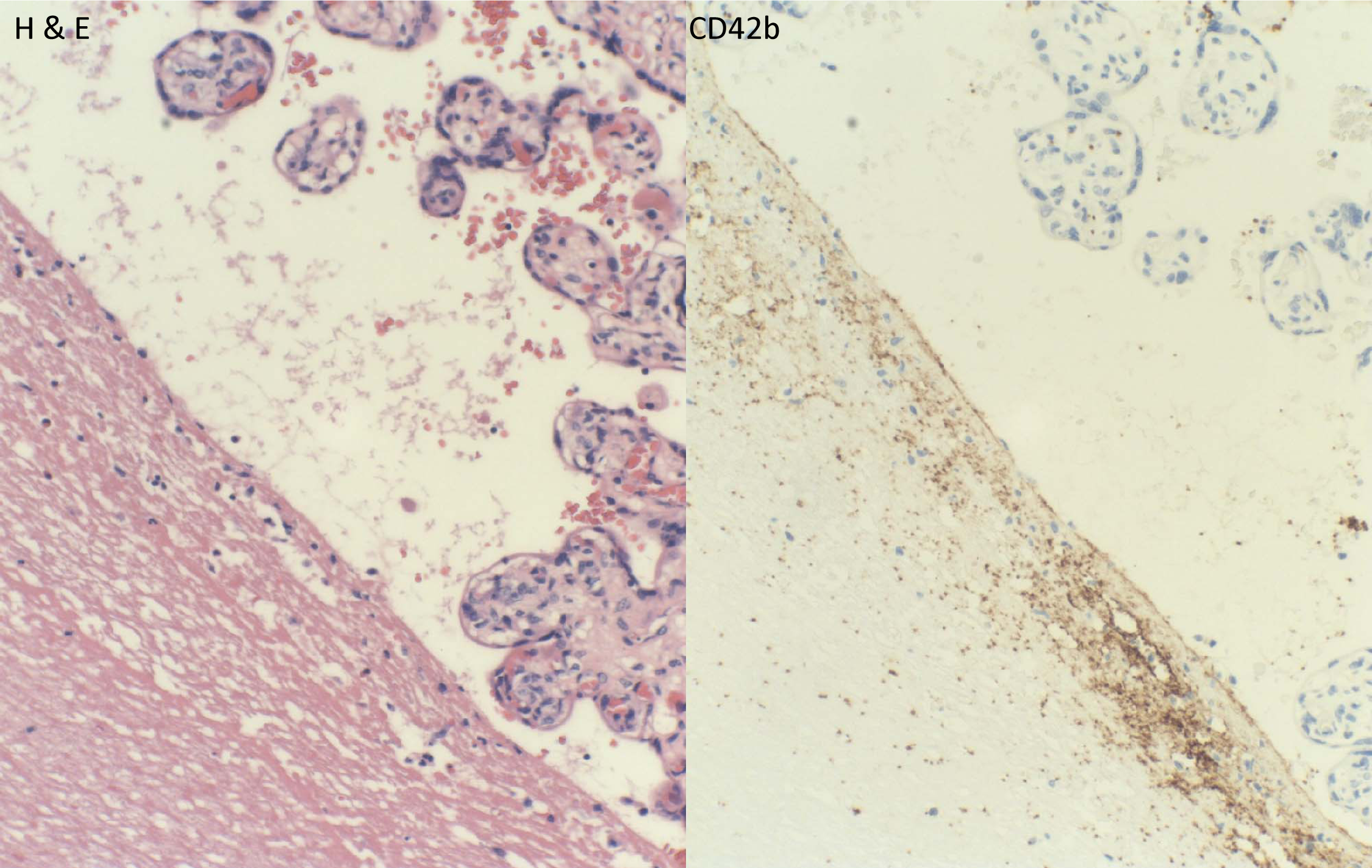
Intervillous thrombus. Intervillous thrombus with H & E stain and the immunohistochemical staining for CD42b.

**Figure 4:**
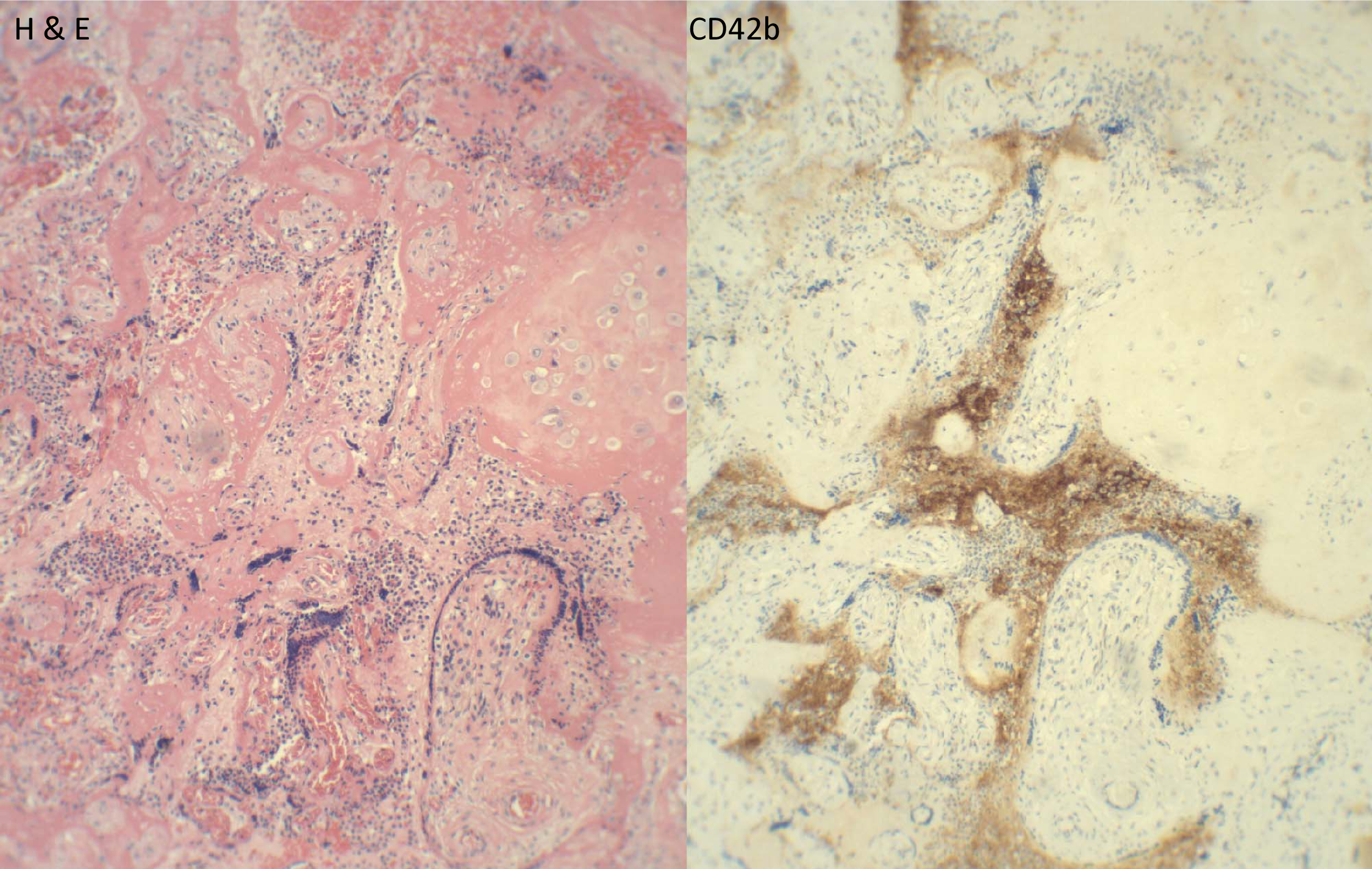
Fibrinoid with chronic villitis. Acute villitis with villous necrosis and peri-villous thrombosis associated with platelet aggregates and increased CD42b immunostaining signals.

**Figure 5:**
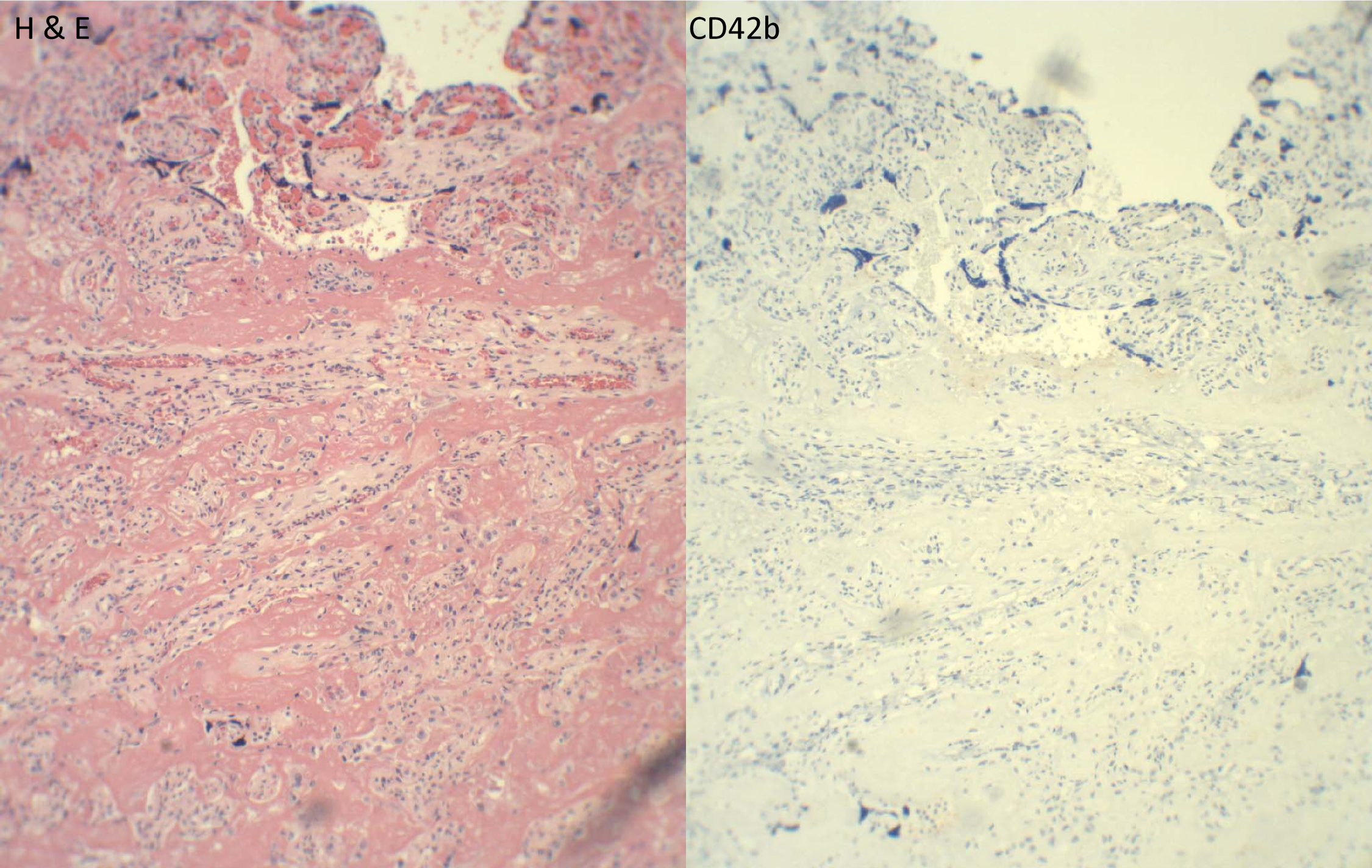
Massive peri-villous fibrinoid deposit (MFI) Maternal floor infarction (massive peri-villous fibrinoid deposit, matrix-type fibrinoid) with no immunohistochemical staining signals for CD42b.

**Figure 6:**
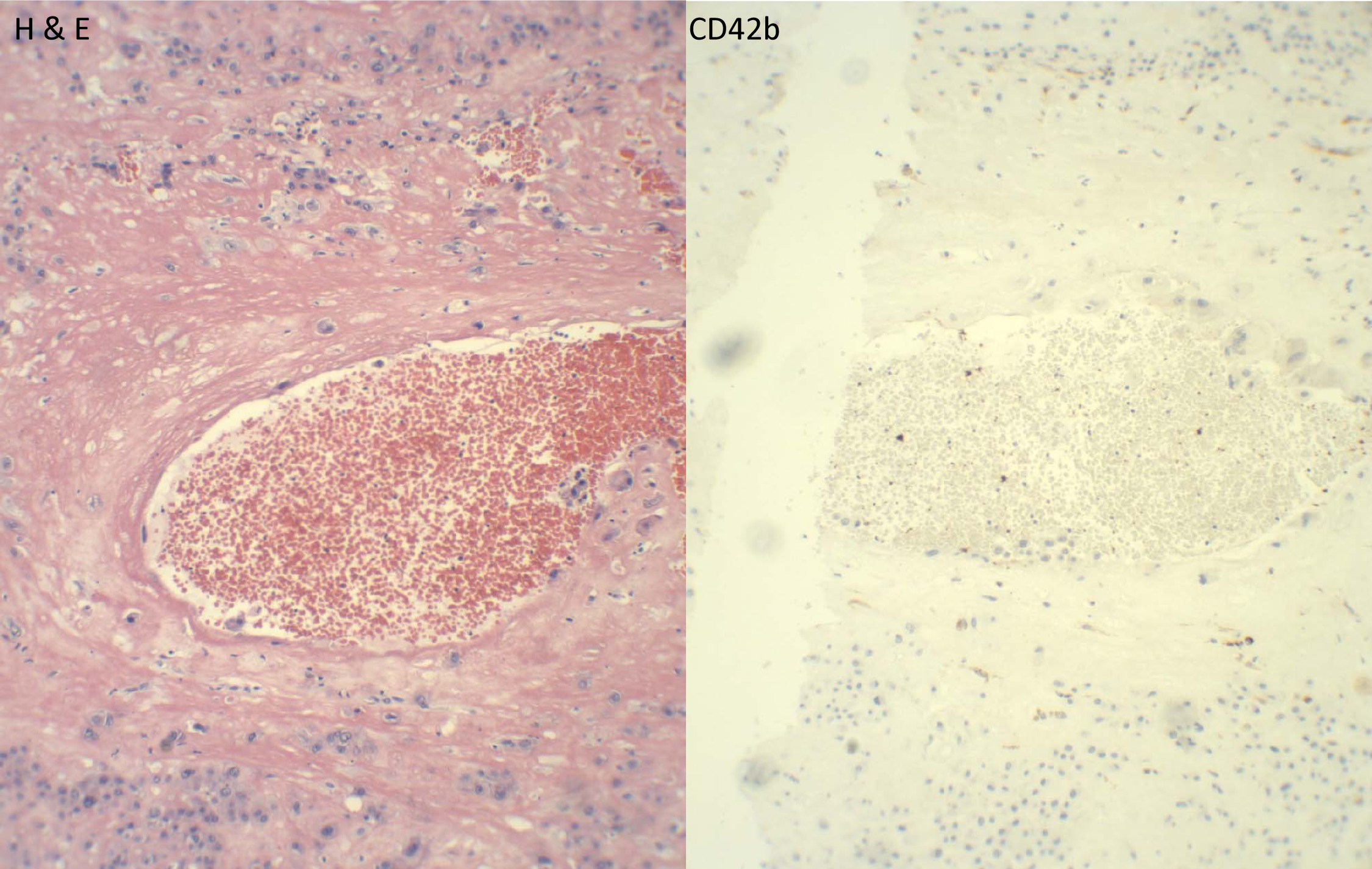
Fibrinoid medial necrosis of decidual vasculopathy. Decidual vasculopathy (fibrinoid medial necrosis) with no immunohistochemical staining signals for CD42b.

## Discussion

Fibrinoid deposit within the placenta is a common finding during routine placental examination, and it mostly does not present as a diagnostic problem for pathologists. In normal pregnancy, fibrinoid at the different anatomic locations may have different functions. Fibrin-type fibrinoid at the subchorionic and the basal region is likely related to the directional change of blood flow, as the normal blood flow within the vessels is laminar in nature, and it becomes turbulent at the vascular bifurcations with changes of direction [14]. The lining cells within these specific locations (syncytiotrophoblasts) can be damaged in a similar fashion to endothelial cell lining at the bifurcation of arteries, leading to thrombosis and platelet aggregates. Similar characteristics can be seen in cases of placental lesions such as accelerated villous maturation in preeclampsia, intervillous thrombosis, infectious villitis or other causes of villous and syncytiotrophoblastic damage, leading to thrombosis. The fibrin-type fibrinoid can be identified by using CD42b immunostaining to highlight the platelet aggregates in difficult cases.

Matrix-type fibrinoid is more complex with various matrix proteins secreted by the trophoblasts in response to various molecular signals [2]. The matrix-type fibrinoid is prominent in maternal floor infarction (MFI/MPFD), and the nature of these fibrinoids in MFI/MPFD is likely related to eosinophilic major basic protein [15-19]. In trophoblastic cysts within the placental tissue (X-cell cyst) the proteinaceous homogenous cystic content was demonstrated to be the major basic protein (MBP) identical to that of eosinophilic granules [15-19]. The function of MBP in placental development is yet to be convincingly demonstrated. Multiple other proteins are identified within the matrix-type fibrinoid with various clinical complications, and these proteins cannot be easily identified during routine clinical practice [2, 5]. Practically the current study is to demonstrate the utility of CD42b immunostaining to distinguish the fibrin-type fibrinoid from the matrix-type fibrinoid during the routine placental examination, and this is especially important in diagnosis of maternal floor infarction (MFI/MPFD).

The clinical significance and functions of fibrinoid within the placenta are largely unknown, and the fibrin-type fibrinoid within the intervillous spaces was initially thought to represent the barrier from the fetal hemorrhage into the maternal circulation in eryhtrobastosis fetalis, and this was supported by the presence of the nucleated red blood cells within the fibrionoid material (Kline hemorrhage) [20]. Subsequent studies support the view that the lining syncytiotrophoblasts of the villous tree depend on the maternal circulation and the damage of the syncytiotrophoblasts leads to intervillous thrombosis [21]. Maternal circulation within the placenta is sinusoidal, and it is different from that of vascular flow [22]. Formation of fibrinoids within the placenta requires more studies.

## Reference

1. Benirschke, K., G.J. Burton, and R.N. Baergen Pathology of the Human Placenta. 6th ed. 2012: Springer.

2. Kaufmann, P., B. Huppertz, and H.G. Frank, The fibrinoids of the human placenta: origin, composition and functional relevance. Ann Anat, 1996. 178(6): p. 485–501.

3. Frank, H.G., et al., Immunohistochemistry of two different types of placental fibrinoid. Acta Anat (Basel), 1994. 150(1): p. 55–68.

4. Khong, T.Y., et al., Sampling and Definitions of Placental Lesions: Amsterdam Placental Workshop Group Consensus Statement. Arch Pathol Lab Med, 2016. 140(7): p. 698–713.

5. Huppertz, B., et al., Extracellular matrix components of the placental extravillous trophoblast: immunocytochemistry and ultrastructural distribution. Histochem Cell Biol, 1996. 106(3): p. 291–301.

6. Faye-Petersen, O.M. and L.M. Ernst, Maternal Floor Infarction and Massive Perivillous Fibrin Deposition. Surg Pathol Clin, 2013. 6(1): p. 101–14.

7. Katzman, P.J. and D.R. Genest, Maternal floor infarction and massive perivillous fibrin deposition: histological definitions, association with intrauterine fetal growth restriction, and risk of recurrence. Pediatr Dev Pathol, 2002. 5(2): p. 159–64.

8. Clemetson, K.J. and J.M. Clemetson, Platelet GPIb-V-IX complex. Structure, function, physiology, and pathology. Semin Thromb Hemost, 1995. 21(2): p. 130–6.

9. Liang, H.P., et al., A common ancestral glycoprotein (GP) 9 1828A>G (Asn45Ser) gene mutation occurring in European families from Australia and Northern Europe with Bernard-Soulier Syndrome (BSS). Thromb Haemost, 2005. 94(3): p. 599–605.

10. Clemetson, J.M., et al., Variant Bernard-Soulier syndrome associated with a homozygous mutation in the leucine-rich domain of glycoprotein IX. Blood, 1994. 84(4): p. 1124–31.

11. Fuchs, B., et al., Distinct role of von Willebrand factor triplet bands in glycoprotein Ib-dependent platelet adhesion and thrombus formation under flow. Semin Thromb Hemost, 2013. 39(3): p. 306–14.

12. Clemetson, K.J. and J.M. Clemetson, Platelet GPIb complex as a target for antithrombotic drug development. Thromb Haemost, 2008. 99(3): p. 473–9.

13. Naeye, R.L., Maternal floor infarction. Hum Pathol, 1985. 16(8): p. 823–8.

14. Cosemans, J.M., et al., The effects of arterial flow on platelet activation, thrombus growth, and stabilization. Cardiovasc Res, 2013. 99(2): p. 342–52.

15. Maddox, D.E., et al., Localization of a molecule immunochemically similar to eosinophil major basic protein in human placenta. J Exp Med, 1984. 160(1): p. 29–41.

16. Vernof, K.K., et al., Maternal floor infarction: relationship to X cells, major basic protein, and adverse perinatal outcome. Am J Obstet Gynecol, 1992. 167(5): p. 1355–63.

17. Wasmoen, T.L., K. Benirschke, and G.J. Gleich, Demonstration of immunoreactive eosinophil granule major basic protein in the plasma and placentae of non-human primates. Placenta, 1987. 8(3): p. 283–92.

18. Wasmoen, T.L., et al., Evidence of eosinophil granule major basic protein in human placenta. J Exp Med, 1989. 170(6): p. 2051–63.

19. Wasmoen, T.L., et al., Association of immunoreactive eosinophil major basic protein with placental septa and cysts. Am J Obstet Gynecol, 1991. 165(2): p. 416–20.

20. Kline and B.S., Microscopic observations of the placental barrier in transplacental erythrocytotoxic anemia (erythroblastosis fetalis) and in normal pregnancy. American Journal of Obstetrics & Gynecology, 1948. 56(2): p. 226--237.

21. Huppertz, B., et al., Villous cytotrophoblast regulation of the syncytial apoptotic cascade in the human placenta. Histochem Cell Biol, 1998. 110(5): p. 495–508.

22. Reynolds, L.P., et al., Evidence for altered placental blood flow and vascularity in compromised pregnancies. J Physiol, 2006. 572(Pt 1): p. 51–8.

